# An international report on bacterial communities in esophageal squamous cell carcinoma

**DOI:** 10.1101/2021.09.29.462325

**Authors:** Jason Nomburg, Susan Bullman, Dariush Nasrollahzadeh, Eric A. Collisson, Behnoush Abedi-Ardekani, Larry O. Akoko, Joshua R. Atkins, Geoffrey C. Buckle, Satish Gopal, Nan Hu, Bongani Kaimila, Masoud Khoshnia, Reza Malekzadeh, Diana Menya, Blandina T. Mmbaga, Sarah Moody, Gift Mulima, Beatrice P. Mushi, Julius Mwaiselage, Ally Mwanga, Yulia Newton, Dianna L. Ng, Amie Radenbaugh, Deogratias S. Rwakatema, Msiba Selekwa, Joachim Schüz, Philip R. Taylor, Charles Vaske, Alisa Goldstein, Michael R. Stratton, Valerie McCormack, Paul Brennan, James A. DeCaprio, Matthew Meyerson, Elia J. Mmbaga, Katherine Van Loon

## Abstract

The incidence of esophageal squamous cell carcinoma (ESCC) is disproportionately high in the eastern corridor of Africa and parts of Asia. Emerging research has identified a potential association between poor oral health and ESCC. One proposed biological pathway linking poor oral health and ESCC involves the alteration of the microbiome. Thus, we performed an integrated analysis of four independent sequencing efforts of ESCC tumors from patients from high- and low-incidence regions of the world. Using whole genome sequencing (WGS) and RNA sequencing (RNAseq) of ESCC tumors and WGS of synchronous collections of saliva specimens from 61 patients in Tanzania, we identified a community of bacteria, including members of the genera *Fusobacterium, Selenomonas, Prevotella, Streptococcus, Porphyromonas, Veillonella*, and *Campylobacter*, present at high abundance in ESCC tumors. We then characterized the microbiome of 238 ESCC tumor specimens collected in two additional independent sequencing efforts consisting of patients from other high-ESCC incidence regions (Tanzania, Malawi, Kenya, Iran, China). This analysis revealed a similar tumor enrichment of the ESCC-associated bacterial community in these cancers. Because these genera are traditionally considered members of the oral microbiota, we explored if there is a relationship between the synchronous saliva and tumor microbiomes of ESCC patients in Tanzania. Comparative analyses revealed that paired saliva and tumor microbiomes are significantly similar with a specific enrichment of *Fusobacterium* and *Prevotella* in the tumor microbiome. Together, these data indicate that cancer-associated oral bacteria are associated with ESCC tumors at the time of diagnosis and support a model in which oral bacteria are present in high abundance in both saliva and tumors of ESCC patients. Longitudinal studies of the pre-diagnostic oral microbiome are needed to investigate whether these cross-sectional similarities reflect temporal associations.

## INTRODUCTION

Esophageal cancer is the sixth most common cause of cancer-related death worldwide (1).There are two histologic subtypes of esophageal cancer with distinct biological characteristics, geographic distributions, and risk factors (2). Esophageal adenocarcinoma is the most common histologic form of esophageal cancer in high-income countries and is associated with factors including gastroesophageal reflux disease, Barrett’s esophagus, and obesity (3, 4). By contrast, esophageal squamous cell carcinoma (ESCC) represents more than 90% of worldwide esophageal cancer cases and is the dominant histology in low-resource settings. In particular, there are two main regions where ESCC is endemic: (1) the Asian esophageal cancer belt, extending from western/northern China to central and southeast Asia; and (2) the eastern corridor of Africa, extending from Ethiopia to South Africa (5, 6).

Emerging research has identified a possible association between poor oral health and ESCC. Studies from Asia, Europe, Latin America, Kenya, and Iran have reported associations of ESCC with poor oral hygiene, chronic periodontal disease, dental decay, and tooth loss (7-16). Recently, three parallel case-control studies in Kenya and Tanzania, conducted as part of the African Esophageal Cancer Consortium (AfrECC) and ESCCAPE (esccape.iarc.fr) collaborations, reported possible associations of poor or infrequent oral hygiene with increased risk for ESCC in East Africa (17-20).

Alterations of the oral microbiome due to poor oral health is one proposed biological pathway that could explain the link between oral health and ESCC. Many bacterial genera associated with gastrointestinal cancers contain species that are traditionally associated with healthy or diseased oral microbiomes. For example, *Helicobacter pylori* was discovered to be associated with gastric cancers and mucosa-associated lymphoid tissue (MALT) lymphomas, indirectly by promoting gastric inflammation and directly by influencing cellular signaling (21). Similarly, bacteria of the genera *Fusobacterium, Selenomonas*, and *Prevotella* are enriched in colorectal cancers (22-24) and can be visualized invasively within tumor tissue (25). *Fusobacterium*, in particular, has been reported to promote carcinogenesis through the selective expansion or inhibition of certain classes of immune cells (26) and may drive cellular proliferation by stimulating Wnt/β-catenin signaling (27, 28). Other bacterial genera such as *Porphyromonas, Campylobacter*, and *Streptococcus* have emerging associations with various human gastrointestinal cancers (29-35).

As part of ongoing investigation into the microbiome’s association with ESCC, we performed an integrated analysis of four independent sequencing efforts including ESCC tumors from patients from both high- and low-incidence regions of the world. In addition, we investigated the relationship between the microbiomes of matched ESCC tumors and saliva specimens in a subset of ESCC cases.

## RESULTS

### Study Population

To evaluate the potential role of the host microbiota in ESCC, we investigated the microbiome of 299 ESCC specimens from patients in five different countries with a high incidence of ESCC. Specimens were collected through four independent sequencing efforts (**Figure 1A**). Specimens consisted of whole genome sequencing (WGS) and RNA sequencing (RNAseq) data from the tumor and saliva of 61 patients from Tanzania (the “MUHAS Tanzania” cohort) (36), RNAseq data from the tumors of 30 ESCC patients in Malawi (the “UNC Project – Malawi” cohort) (37), and WGS from 208 additional samples of tumors from patients in high ESCC incidence regions, including specimens from ESCC patients in Tanzania (n=18) and Kenya (n=64) that were collected in the ESCCAPE studies (esccape.iarc.fr) and specimens from ESCC patients in East Golestan, Iran (n=55) and Shanxi, China (n=71) that were sequenced as part of the Cancer Research UK Mutographs project (“Mutographs” cohorts) (38). In addition, we analyzed WGS data of ESCC from The Cancer Genome Atlas (39), which includes a small number of tumors from patients in low-incidence geographic regions including the United States (n=3), Ukraine (n=3), Vietnam (n=22), and Russia (n=8) (the “TCGA” cohort). Patient characteristics are shown in Table 1.

**Figure 1.**
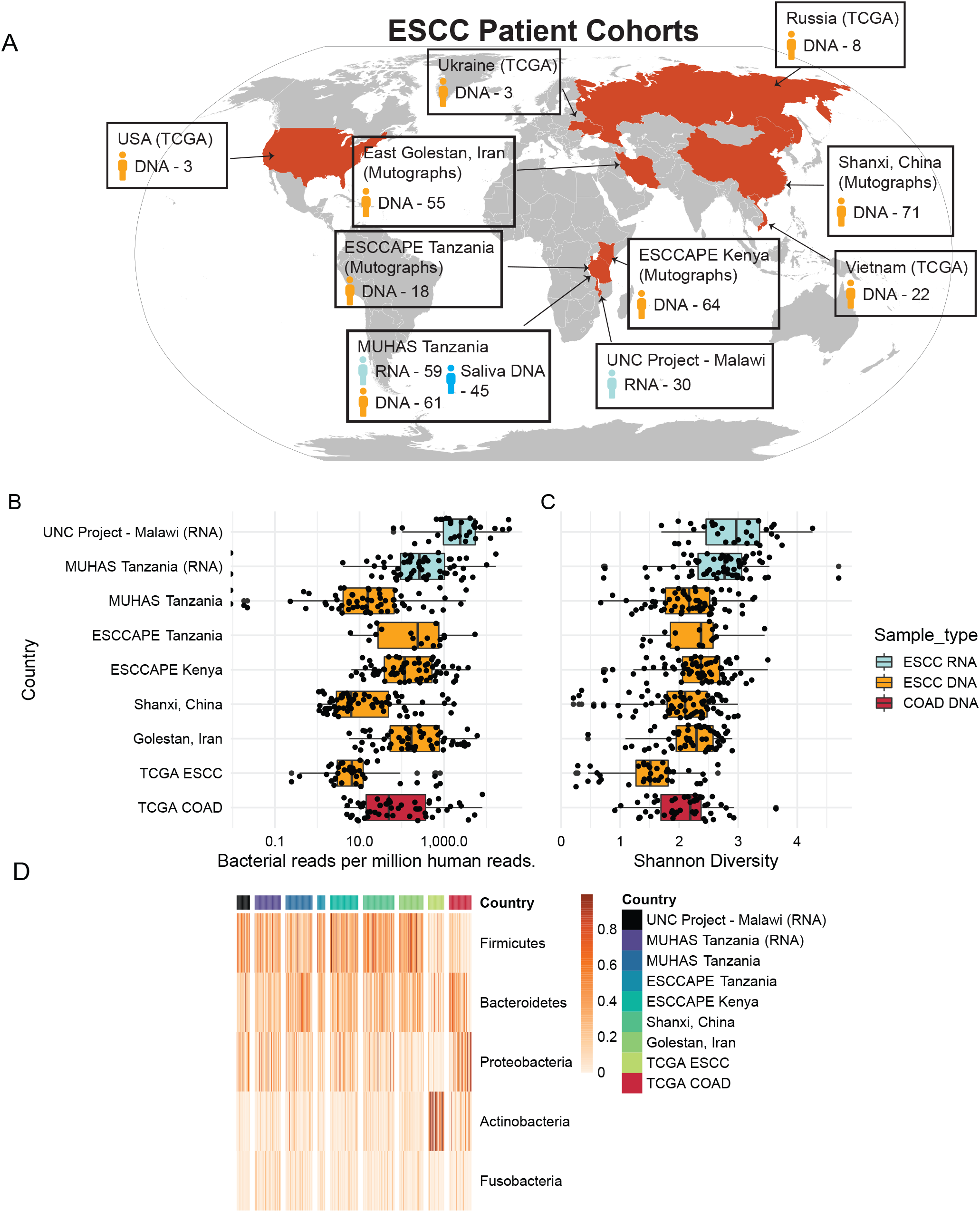
Microbiome structure and composition of ESCC tumors. A. Description of ESCC patients, and sample types, assessed in this study. TCGA – The Cancer Genome Atlas; ESCCAPE – Esophageal Squamous Cell Carcinoma African Prevention Research; Mutographs – Cancer Research UK Mutographs Project. B. Bacterial burden of ESCC tumors for each patient cohort. Units are bacterial reads per million human reads as determined by GATK-PathSeq analysis. Each dot represents one sample. Analyte type (RNA or DNA) and tumor type (ESCC or COAD) are indicated by color. C. Shannon diversity of ESCC tumors for each patient cohort. Shannon diversity was determined for each sample at the genus level based on genera that are at least 1% relative abundance. Each dot represents one sample. Analyte type (RNA or DNA) and tumor type (ESCC or COAD) are indicated by color. D. Heatmap describing the relative abundance of the five top phyla sorted by average phylum relative abundance. Each column represents one sample. Rows represent the indicated phyla. Units are relative abundance. Samples from each cohort are WGS unless noted with “(RNA)”, in which case they are RNAseq.

**TABLE 1.**

| **TABLE 1** |  |  |  |  |  |  |
| --- | --- | --- | --- | --- | --- | --- |
| **Study** | **Tanzania** | **Malawi\*\*** | **ESCAPE Tanzania\*\*\*** | **ESCAPE Kenya\*\*\*** | **East Golestan, Iran\*\*\*\*** | **Shanxi, China\*\*\*\*** |
| No. cases included | 61 | 30 | 18 | 65 | 55 | 71 |
| **Demographics** |  |  |  |  |  |  |
| Median age (IQR) | 49 (44-62) | 56 | 65 (61-73) | 64 (53, 71) | 62 (54,73) | 56 (50, 64) |
| % male | 67% | 45.8% | 61% | 68% | 55% | 56% |
| **Status at diagnosis** |  |  |  |  |  |  |
| Weight (kg), median (IQR) |  |  | 44 (40-52) | 52 (46, 60) |  |  |
| Body mass index (kg/m<sup>2</sup>) median (IQR) |  |  | 15.8 (15.4, 19.1) | 19.5 (15.6, 22.0) |  |  |
| Median months ill before coming to endoscopy (IQR) |  |  | 2 (1, 6) | 3 (2, 4.5) |  |  |
| HIV status: |  |  |  |  |  |  |
| Positive | 2 (3.2%) | 10 (16.9%) | 1 (5%) | 5 (8%) |  |  |
| Negative | 36 (59.0%) | 44 (74.6%) | 10 (56%) | 48 (74%) |  |  |
| Not known | 23 (37.7%) | 5 (8.5%) | 7 (39%) | 12 (18%) |  |  |
| **Key lifestyle habits** |  |  |  |  |  |  |
| N (%) ever tobacco users |  |  | 11 (61%) | 38 (58%) | 17 (31%) | 35 (49%) |
| N (%) who brush teeth daily: |  |  |  |  |  |  |
| With toothbrush |  |  | 12 (67%) | 16 (25%)* |  |  |
| With stick |  |  | 6 (33%) | 10 (15%) |  |  |
| Median no missing teeth (IQR) |  |  | 3 (1, 5) | 4 (1, 8) |  |  |

\*N=22 (34%) brush once per week or never, n=17 (26%) brush 2 to 6 times/week
\*\*Indicates demographics are from the entire patient population, consisting of both included and unincluded patients.
\*\*\*Indicates demographic percentages are from the entire patient population, with discrete counts scaled to the number of cases included.
\*\*\*\*Indicates demographic information is exclusively for included patients.

### Bacterial populations are abundant and diverse in ESCC tumors

We used the metagenomic analysis tool GATK-PathSeq (40) to process the RNAseq and WGS data. GATK-PathSeq uses a sequential mapping strategy to assign reads to human and microbial reference genomes, resulting in detailed information on sequencing reads of human and microbial origin (**Figure S1A**). We likewise used GATK-PathSeq to process WGS data sets from 50 colon adenocarcinoma (COAD) specimens available from TCGA (41) for comparison, as there is strong evidence of microbial associations with COAD (22-25).

The bacterial burden of ESCC tumors ranged from 10 to 1000 bacterial reads per million human reads, similar to numbers observed in TCGA COAD (**Figure 1B**). Furthermore, the Shannon diversity of bacterial populations at the genus level ranged from 2 to 3 (**Figure 1C**). By comparison, ESCC-associated bacterial communities are as diverse or more diverse than TCGA COAD. At the phylum level, ESCC bacterial populations generally consist of *Firmicutes, Bacteroidetes, Proteobacteria, Actinobacteria*, and *Fusobacteria* (**Figure 1D, Figure S1B**). Of note, the higher than expected abundance of the phylum *Actinobacteria* specifically in the TCGA ESCC samples is attributable, in particular, to a very high abundance of the genus *Tetrasphaera* (**Figure S1C**). This is evidenced by a depressed Shannon diversity of *Actinobacteria* genera in these samples (**Figure S1D**) and may indicate contamination of the TCGA ESCC samples. *Actinobacteria* have been reported as a source of contaminating reads in TCGA gastrointestinal cancer samples (42).

### Bacterial genera associated with carcinogenesis are observed at high relative abundance in ESCC tumors from Tanzania

To determine if bacteria with known associations with cancer are present in ESCC, we first analyzed the sequencing series of the 61 ESCC cases from the MUHAS Tanzania cohort with both WGS and RNAseq data. The paired WGS and RNAseq data from these tumors allowed investigation of bacterial communities at the DNA and RNA levels. Both WGS and RNAseq data revealed high relative abundance of bacterial genera previously associated with carcinogenesis in these ESCC tumors (**Figure 2A, 2B**). The high relative abundance of the *Fusobacterium* genus was particularly notable. Other bacterial genera of interest include *Streptococcus, Porphyromonas, Campylobacter, Prevotella, Veillonella*, and *Selenomonas*, many of which have been associated with gastrointestinal malignancies alongside or independently of *Fusobacterium* (25, 29, 32, 34, 43). The mean Jaccard similarity index between tumor RNAseq and WGS data from the same tumor is 0.54, greater than the average Jaccard similarity index of random RNAseq-WGS pairs (0.36), indicating that bacterial populations inferred from WGS and RNAseq data are generally consistent (**Figure 2C**).

**Figure 2.**
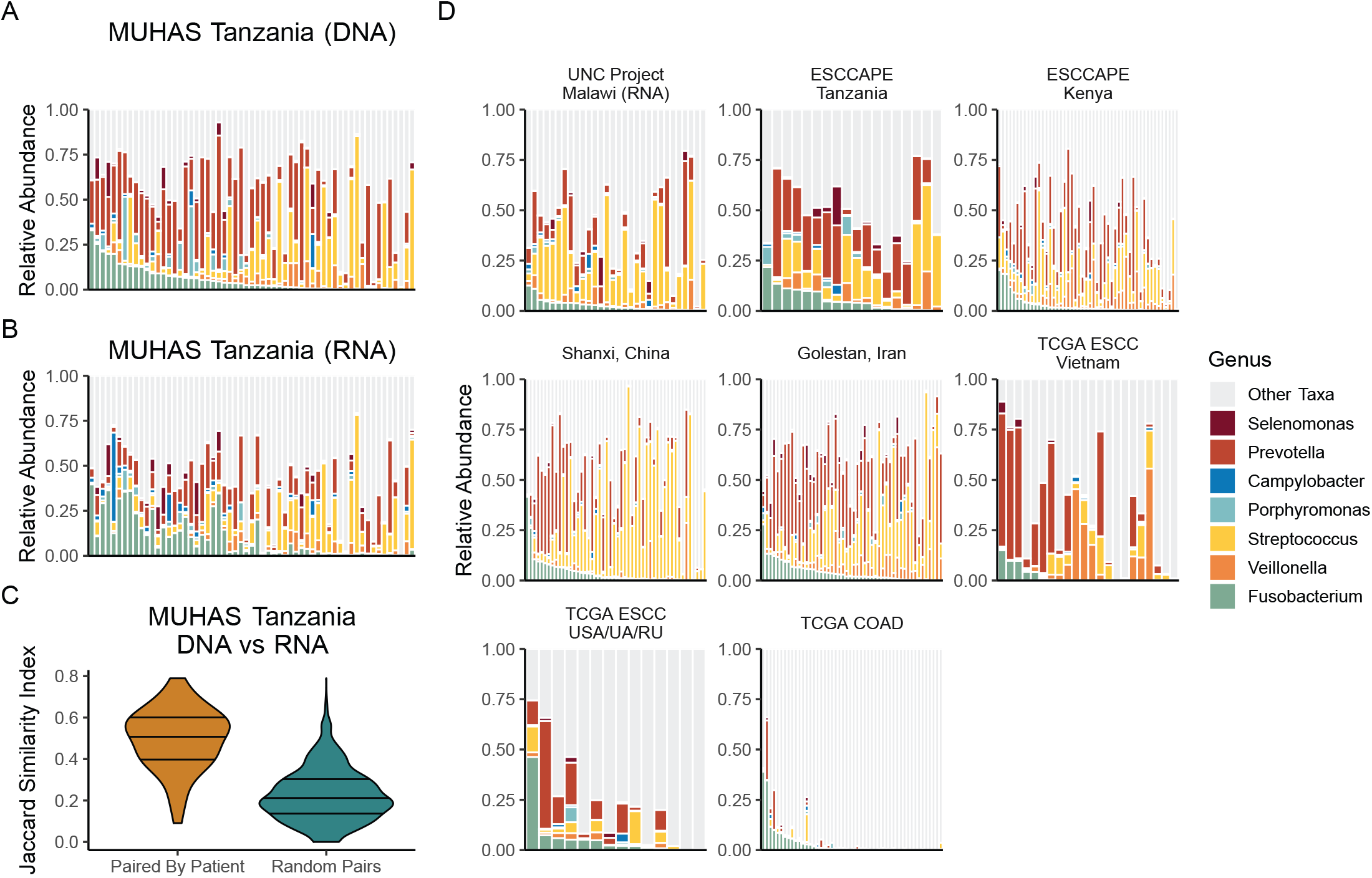
Identification of bacterial genera associated with carcinogenesis. A. Bacterial genera relative abundance of WGS data from the MUHAS Tanzania cohort. Each column represents a single sample. Samples are ordered by decreasing Fusobacterium relative abundance. Units are relative abundance of bacterial genus-mapping reads. Color indicates the genus, and seven genera are specified. Only patients with GATK-PathSeq analysis from both RNAseq and WGS tumor data are plotted (n=59). Columns are ordered by decreasing relative abundance of *Fusobacterium* genus reads. B. Bacterial genera relative abundance of RNAseq data from the MUHAS Tanzania cohort. Each column represents a single sample. Here, column order is dictated according to the patient order in Figure 2A. Units are relative abundance of bacterial genus-mapping reads. Color indicates the genus, and seven genera are specified. Only patients with GATK-PathSeq analysis from both RNAseq and WGS tumor data are plotted (n=59). Samples are ordered in the same order as Figure 2A, which is by *Fusobacterium* genus relative abundance in the WGS data. C. Jaccard index between RNAseq and WGS data of tumors from the MUHAS Tanzania cohort. For the “Paired by Sample” column, Jaccard indices were calculated only between the WGS and RNAseq data from the same tumor (n=59 comparisons). For the “Random Pairs” column, Jaccard indices were calculated between all possible WGS-RNAseq pairs independent of patient of origin to represent the expected random distribution of Jaccard indices (n=3,481 comparisons). Jaccard index was calculated from relative abundance at the genus level based on genera that are at least 1% relative abundance. The width of the violin represents the relative proportion of comparisons with each Jaccard index, and lines indicate 25^th^, 50^th^, and 75^th^ percentiles. D. Bacterial genera relative abundance of the remaining patient cohorts, including RNAseq and WGS data as indicated. Each column represents a single sample. Samples are ordered by decreasing Fusobacterium relative abundance within each patient cohort. Units are relative abundance of bacterial genus-mapping reads. Color indicates the genus, and seven genera are specified. Here, if there were more than 50 samples in a patient cohort, 50 samples were randomly selected for visualization. USA – United States, UA – Ukraine, RU – Russia. All cohorts consist of WGS data, with the exception of the tumors from Malawi which are RNAseq. (Number of samples plotted: UNC Project - Malawi 30; ESCCAPE Tanzania 18; ESCCAPE Kenya 50; Shanxi, China 50; Golestan, Iran 50; TCGA ESCC Vietnam 22; TCGA ESCC USA/UA/RU 14).

Next, we attempted to determine if similar bacterial genera were also present in ESCC from patients in high-incidence countries beyond Tanzania. Investigation of RNA sequencing data from patients in Malawi, WGS data from patients in Kenya, China, and Iran, as well as from the independent ESCCAPE Tanzania patient group revealed pervasive evidence of similar bacterial genera in the tumors of these patients (**Figure 2D, Figure S2A**). To investigate if similar microorganisms were found in ESCC tumors from patients in low-incidence regions, we investigated WGS data from ESCC tumors originating from USA, Ukraine, Vietnam, and Russia that were available through TCGA. While the number of samples available from low-incidence regions is low and relies on a single sequencing effort, we found that the tumors of many of these patients contain similar bacterial genera (**Figure 2D, Figure S2A)**. Colon cancers from the TCGA COAD cohort revealed evidence of *Fusobacterium*, as expected; however, these COAD samples were notable for much lower relative abundance of the other genera of interest, when compared to ESCC tumors.

### Evaluation of association between saliva and tumor microbiomes in ESCC patients from Tanzania

We next investigated the similarity between the saliva and tumor microbiomes of ESCC patients. Paired saliva samples were only available from patients in the MUHAS Tanzania cohort (N=45); these paired saliva specimens were analyzed to evaluate bacterial abundance as a proxy for the oral microbiome.

We first assessed the similarity between paired saliva and tumor microbiomes with the Bray Curtis similarity index (44). To avoid potential confounding due to low bacterial read counts in some tumor samples, we limited these analyses to the 21 tumor-saliva pairs that contain appreciable microbial sequencing depth (at least 10,000 bacterial reads each). We found that the saliva and tumor microbiomes from the same patient in the Tanzanian samples are significantly more similar than random saliva-tumor pairs (p=0.0003, Wilcoxon rank sum test) (**Figure 3A**). Next, we asked if there are bacterial genera whose relative abundance in the saliva correlates with their relative abundance in the tumor. For this analysis, we included only common-abundant bacterial genera with at least 1% relative abundance in at least three tumor-oral pairs. The relative abundance of four bacterial genera (*Fusobacterium, Veillonella, Streptococcus*, and *Porphyromonas*) are strongly correlated between tumor and saliva microbiomes, while other common-abundant bacterial genera were not (**Figure 3B**). To assess if any bacterial genera are preferentially enriched in the tumor microbiome relative to the saliva microbiome, we next calculated the difference in the relative abundance of the common-abundant bacterial genera between saliva-tumor pairs. Several genera including *Porphyromonas* and *Veillonella* were at higher relative abundance in the saliva, while *Prevotella* and *Fusobacterium* were enriched in the tumor microbiome (**Figure 3C**). Finally, the relative abundance of tumor-associated bacteria including *Fusobacterium, Prevotella, Selenomonas, Veillonella, Streptococcus*, and *Campylobacter* are strikingly similar between the microbiomes of tumor and oral pairs (**Figure 3D**). Altogether, these data support the hypothesis that there is an association between the oral and tumor microbiome of ESCC patients in Tanzania.

**Figure 3.**
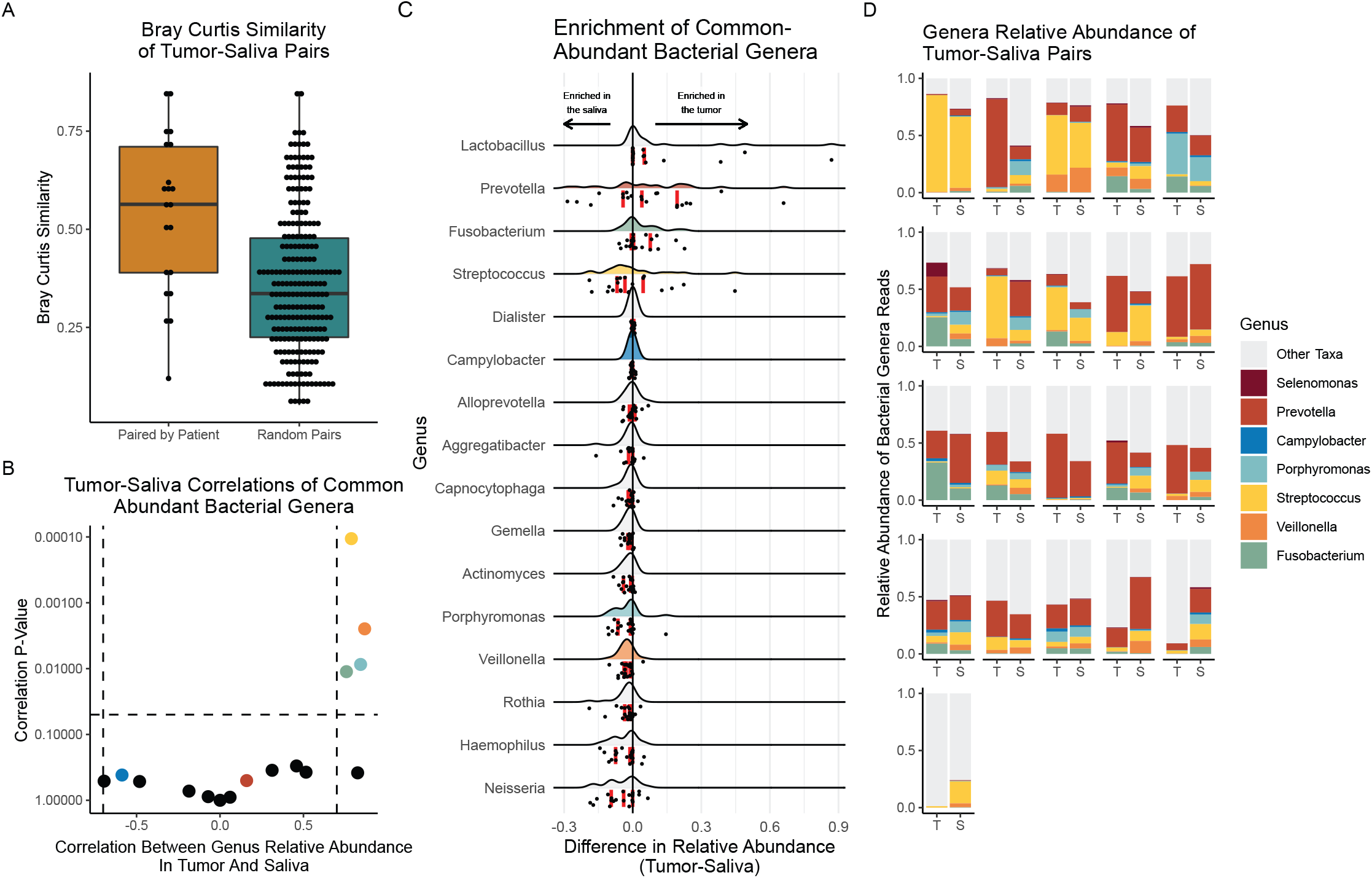
Association between synchronous saliva and tumor microbiomes in Tanzanian ESCC patients. **A**. Bray Curtis Similarity comparing tumor-saliva pairs from patients in the MUHAS Tanzania cohort. Analysis was restricted to the 21 tumor-saliva pairs that contained at least 10,000 bacterial reads. This analysis was conducted at the genus level and using relative abundance. For the “Paired by Patient” column, Bray Curtis Similarity was calculated only between the tumor and saliva WGS data from the same patient. For the “Random Pairs” column, Bray Curtis Similarity was calculated between all possible tumor-saliva pairs independent of patient of origin to represent the expected random distribution of Bray Curtis Similarity. (p=0.0003, Wilcoxon rank sum test). **B**. Correlation between the relative abundance of common-abundant bacterial genera in paired saliva and tumor WGS data. Analysis was restricted to the 21 tumor-saliva pairs that contained at least 10,000 bacterial reads. Common-abundant bacterial genera are bacterial genera that are at least 1% abundance in at least 3 tumor-saliva pairs – 16 bacterial genera made this cutoff. Correlation represents a two-sided Pearson correlation. X-axis is the correlation coefficient, and Y axis is the correlation P-Value plotted on a log scale. **C**. Enrichment of genera in the oral or tumor microbiome. Each row details one of the 16 common-abundant bacterial genera. Each row contains one data point per patient, for a total of 21 data points. The value of each point represents the difference in the relative abundance of the specified genus in the tumor and oral microbiomes of one patient, with positive values indicating a genus is at higher relative abundance in a patient’s tumor. For example, if a genus is at a relative abundance of 0.7 (70%) in the tumor and 0.3 (30%) in the saliva of a patient, the plotted value for that genus and that patient is 0.4. Curves represent the distribution of this relative abundance difference across the tumor-oral pairs, with dots indicating individual tumor-oral pairs. Vertical red lines indicate quartiles. **D**. Relative abundance bar charts of tumor-saliva pairs. Analysis was restricted to the 21 tumor-saliva pairs that contained at least 10,000 bacterial reads. Units are relative abundance of bacterial genus-mapping reads. Color indicates the genus, and seven genera are specified. (abbreviations: T – tumor, S – saliva).

## DISCUSSION

This report provides an analysis of bacterial communities present in ESCC tumors from nine countries from different regions of the world, analyzed in four independent sequencing efforts. We found traditionally oral, cancer-associated, bacterial genera in tumors from patients in Tanzania, Malawi, Kenya, China, and Iran. These results provide evidence that these bacterial genera may be associated with ESCC in high-incidence regions. We also identified similar bacterial genera in ESCC tumors from low-incidence regions, although this finding is based on a small sample size and only one sequencing cohort. Finally, in a sub-analysis of tumor and saliva pairs available from Tanzania, we demonstrated that the synchronous collected saliva and tumor microbiomes of ESCC patients are strikingly similar at the time of diagnosis; in particular, we identified a specific correlation between the saliva and tumor relative abundance of the bacterial genera *Fusobacterium, Veillonella, Streptococcus*, and *Porphyromonas*, with *Prevotella* and *Fusobacterium* significantly enriched in the tumor microbiome.

Many of the bacterial genera identified in this study have been previously implicated in the carcinogenesis of gastrointestinal cancers. For example, studies have found that oral microbiota including *Fusobacterium, Prevotella, Selenomonas, Veillonella, Streptococcus*, and *Campylobacter* can be used to distinguish individuals with colorectal cancer from healthy controls (45), and that *Fusobacterium nucleatum* strains that colonize the oral cavity and tumors of patients with colorectal cancer are identical in some patients (46), raising the possibility that the oral cavity is a source of extra-oral cancer microbiota. Our group has previously shown that *Fusobacterium, Selenomonas*, and *Prevotella* can be visualized invasively within colorectal tumors and liver metastases (25). Fusobactium nucleatum has been previously identified in esophageal cancers and is associated with shorter survival (47). Members of the genus *Porphyromonas* have been previously observed invasively within ESCC tumors (29) and have been reported to promote oral squamous cell carcinoma through a variety of mechanisms (30, 31). *Campylobacter jejuni* has been reported to promote tumorigenesis in mice (32), and *Streptococcus* species have been identified in human esophageal cancers (33). In addition, the striking association of *Streptococcus bovis* with colorectal cancer has led to the recommendation that colonoscopy be performed upon detection of *Streptococcus bovis* bacteremia or endocarditis (34, 35). Oral commensal bacteria such as *Veillonella* species have been previously implicated in pathogenesis of lung cancer (43). A prospective cohort of American patients (48) and a study of Japanese patients (49) likewise found that oral microbiome composition reflects risk of esophageal cancers

We found that bacterial genera including *Fusobacterium, Prevotella, Selenomonas, Veillonella, Streptococcus*, and *Campylobacter* are pervasive in the microbiome of ESCC tumors from patients in high-incidence regions. Moreover, the bacterial composition of ESCC tumors is remarkably similar across countries in those high-incidence regions, raising the possibility that these bacterial genera may be involved in ESCC carcinogenesis or that they may colonize tumors as a result of the common clinical presentation of patients with severe dysphagia. Notably, there are several alternative hypotheses that warrant mention. For example, it is possible that the ESCC-associated bacterial genera simply represent common members of the esophageal microbiome (50) and that the microbial populations we observed in these cancers are not significantly different from those found in normal esophagus tissue. A limitation of our study is a lack of normal esophageal tissue from ESCC cases or healthy controls in these settings, which would allow us to address this possibility. Another possible explanation is that ESCC tumors provide a favorable niche in which these bacteria are sequestered and allowed to colonize due to the propensity of this disease to cause malignant obstruction. Thus, it is plausible that ESCC-associated bacteria are not necessarily promoting ESCC carcinogenesis but rather represent passengers resulting from the sequestration of oral secretions proximal to an obstructing tumor. While the previous association of these bacterial genera with other cancers is consistent with the hypothesis that they influence carcinogenesis of ESCC, future studies are necessary to identify which, if any, direct influences these bacterial genera have upon ESCC carcinogenesis. Nevertheless, even if these bacterial genera do not have a role in increasing ESCC risk, but arise at the time of disease onset, they may have an important role to play as part of a non-invasive early-detection biomarker. Finally, a concern of all microbiome analyses is that observed bacteria can be a consequence of contamination at some step between tumor harvest and sequencing. While some TCGA samples may be contaminated by *Actinobacteria* as previously noted, the presence of *Fusobacterium, Prevotella, Selenomonas, Veillonella, Streptococcus*, and *Campylobacter* in four independently collected cohorts indicates that these finding are unlikely due to contamination.

While this study focused on the presence of bacteria with ESCC in high-incidence regions, we found evidence of similar cancer-associated bacteria in tumors in patients from low-incidence regions (USA, Ukraine, Vietnam, and Russia). A limitation of this assessment is the small sample size (n=36) and reliance on a single TCGA cohort that likely contains contaminants (42). Regardless, this finding does not exclude the possibility that the microbiome could be a factor driving patterns of ESCC incidence. For example, it is possible that the prevalence of ESCC-associated bacteria in people could vary across regions, which in turn could drive these differing rates of ESCC incidence. This is an important topic for future study.

We found that the structure of synchronous paired tumor and oral microbiomes were strikingly similar. It is possible that this similarity is driven by transient contact of saliva and its associated microbiome with the tumor (e.g., during swallowing or tumor extraction). However, we found that only four of sixteen common-abundant bacterial genera correlate in abundance between the tumor and oral microbiomes, suggesting tumor-oral microbiome similarity is not driven exclusively by “in-trans” interactions between the saliva and tumor. We also found that genera including *Prevotella* and *Fusobacterium* are often specifically enriched in the tumor microbiome, supporting a model where specific oral bacterial preferentially colonize the tumor. A caveat of this study is that we infer oral bacterial populations from the saliva, despite diverse communities of bacteria throughout the oral cavity (51). However, we do observe *Fusobacterium* in the saliva despite its general association with periodontal plaques (52), suggesting saliva is capable of detecting periodontal pathogens. Additionally, because the samples studied here are from patients with late-stage disease, it is possible that tumor-induced changes to upper-gastrointestinal physiology and dysphagia symptom-induced major dietary changes could themselves alter the oral microbiomes of these patients. The previous findings from the ESCCAPE studies in Kenya and Tanzania (17, 19) which found strong associations with dental staining (Ors > 10) and for which photographic validation studies suggest that most dental staining was not fluorosis, also point to a recent build-up of chromogenic bacteria. Studies of the oral microbiome of patients at earlier stages of ESCC and in prospective studies are necessary to address this possibility. We restricted our analysis to 21 tumor-oral pairs that have a sufficient number of bacterial reads (at least 10,000). It is likely that excluded samples are not molecularly distinct from included samples but that the relatively low bacterial read counts in some tumors is simply reflective of low sequencing depth.

Our observation of similar tumor and saliva microbiomes in ESCC patients is especially notable considering emerging evidence linking periodontal disease and poor oral health with increased risk of various cancers (17, 53, 54). This raises several important open questions. It will be essential to determine if there is a difference in the oral prevalence of these identified cancer-associated bacteria between ESCC patients and non-patients earlier in the natural history of the disease, for example through comparisons of patients with esophageal squamous dysplasia and healthy controls. Because the prevalence of these bacteria may be associated with factors such as oral health, hygiene, and diet, studies of the impact of these factors on the oral microbiome in the general population would inform whether the oral microbiome is on a pathway linking oral hygiene to ESCC risk and may have a role in prevention.

In conclusion, we show that cancer-associated, traditionally-oral bacteria including the genera *Fusobacterium, Selenomonas, Prevotella, Streptococcus, Porphyromonas, Veillonella*, and *Campylobacter* are highly abundant within ESCC tumors from patients in high-ESCC incidence regions. We also show that there is a correlation between the genus composition of the saliva microbiome and the ESCC tumor microbiome of some ESCC patients. These findings will be foundational for future studies to understand if and how bacteria influence ESCC pathogenesis and to understand the role of the oral microbiome in this process. Finally, this study highlights the benefit of collaborative investigation to evaluate the international heterogeneity of this disease.

## MATERIALS AND METHODS

### Sample acquisition and sequencing

The sample acquisition and sequencing methods for the studies from the MUHAS Tanzania cohort (n=61) (36) and UNC Project - Malawi cohort (n=30) (11) have been previously described. Samples sequenced in the Mutographs study (n=210) (38) originated from patients in Golestan, Iran (n=55), ESCCAPE case-control studies in Tanzania (n=18) (19) and Kenya (n=64) (17), and patients in Shanxi, China (n=71). TCGA ESCC (n=36) and COAD samples (n=51) have been previously described (39, 41). The TCGA ESCC cohort includes tumors from patients in United States (n=3), Ukraine (n=3), Vietnam (n=22), and Russia (n=8), regions which have lower incidence of ESCC.

### Metagenomic analysis

GATK-PathSeq (40) was used to conduct computational subtraction of human-mapping reads from input RNAseq and WGS datasets. GATK-PathSeq works by first mapping reads to a host reference database consisting of the human genome grch38 and various supplemental human reference sequences. Next, non-human reads are mapped against a comprehensive microbial database, and microbe read assignments are reported for further study. From the MUHAS Tanzania cohort, a total of 61 tumor WGS samples, 45 saliva WGS samples, and 59 RNAseq samples were processed through GATK-PathSeq.

Bacterial abundance analyses and plotting were conducted in R (v3.5.1). To calculate relative abundance at a phylogenetic level (e.g., phylum or genus), GATK-PathSeq results were filtered for taxa at the level, and relative abundance was calculated for each taxon as follows: (# of taxon reads)/(total # reads at the selected phylogenetic level). The rows of all bacterial abundance heatmaps are arranged according to the mean abundance across all samples. The sample order of relative abundance stacked barplots were determined based on *Fusobacterium* genus relative abundance except where noted. In **Figure 2D**, if any cohort contained more than 50 samples, 50 samples were randomly selected for plotting. The distribution of relative abundances of genera of interest in all samples can be found in **Figure S2**, where width of each violin represents the relative distribution of observed bacterial relative abundance for all patients in each patient cohort.

Jaccard distance between RNAseq and WGS data from each ESCC tumor was calculated in R based on bacterial genera with at least 1% relative abundance. The qualitative Jaccard index was used in this case because the comparison was between DNA and RNA analytes which would not be expected to be quantitatively identical.

### Tumor-saliva similarity

Only tumor-saliva pairs from the MUHAS Tanzania cohort with at least 10,000 reads mapped to the bacterial superkingdom were available for analysis. This resulted in a total of 21 tumor-oral pairs. Bray-Curtis dissimilarity metrics between tumor-oral pairs were calculated using the R package vegan (55). **Figure 3A** presents the Bray-Curtis *similarity* (1 – Bray-Curtis dissimilarity), for each tumor-oral pair.

To determine the correlation between the relative abundance of specific genera between tumor and saliva microbiomes, common-abundant genera that are at least 1% abundance in at least 3 tumor-oral pairs were identified. This resulted in the identification of 16 common-abundant genera. Correlations represent a two-sided Pearson correlation coefficient. To determine tumor-oral enrichment of common-abundant genera, the difference in relative abundance of each genus between each tumor-oral pair was plotted (**Figure 3C**). For the relative abundance bar plots of tumor-saliva pairs (**Figure 3D**), bacterial genera that had been highlighted in previous figures are labeled.

### Code and processed data availability

All GATK-PathSeq output files and reproducible analysis and plotting R Notebooks are available.

Zenodo: https://doi.org/10.5281/zenodo.4750577

GitHub: https://github.com/jnoms/ESCC_microbiome

Furthermore, all analysis and figures can be automatically reproduced through a series of Google Colab documents.

Figure 1 and Supplementary Figure 1: https://github.com/jnoms/ESCC_microbiome/blob/main/collab/Figure1.ipynb

Figure 2 and Supplementary Figure 2: https://github.com/jnoms/ESCC_microbiome/blob/main/collab/Figure2.ipynb

Figure 3. : https://github.com/jnoms/ESCC_microbiome/blob/main/collab/Figure3.ipynb

## ABBREVIATIONS

AFRECC: African Esophageal Cancer Consortium
COAD: Colon adenocarcinoma
ESCA: Esophageal adenocarcinoma
ESCC: Esophageal squamous cell carcinoma
ESCCAPE: Esophageal Squamous Cell Carcinoma African Prevention Research
MUHAS: Muhimbili University of Health and Allied Sciences
RNAseq: RNA sequencing
TCGA: The Cancer Genome Atlas
WGS: Whole genome sequencing

## ACKNOWLEDGMENTS

We thank Liu et al. (37) for providing access to sequencing data from ESCC patients in Malawi (dbGaP accession phs001448.v1.p1). Portions of this research were conducted using the O2 High Performance Computing Cluster, supported by the Research Computing Group at Harvard Medical School. We thank Aleksandar Kostic for his helpful discussions. We thank the teams from MUHAS-ORCI-UCSF Cancer Collaboration, UNC-Malawi Project, Mutographs, ESCCAPE, and TCGA for specimen and data collection.

## FIGURE LEGENDS

**Figure S1.**
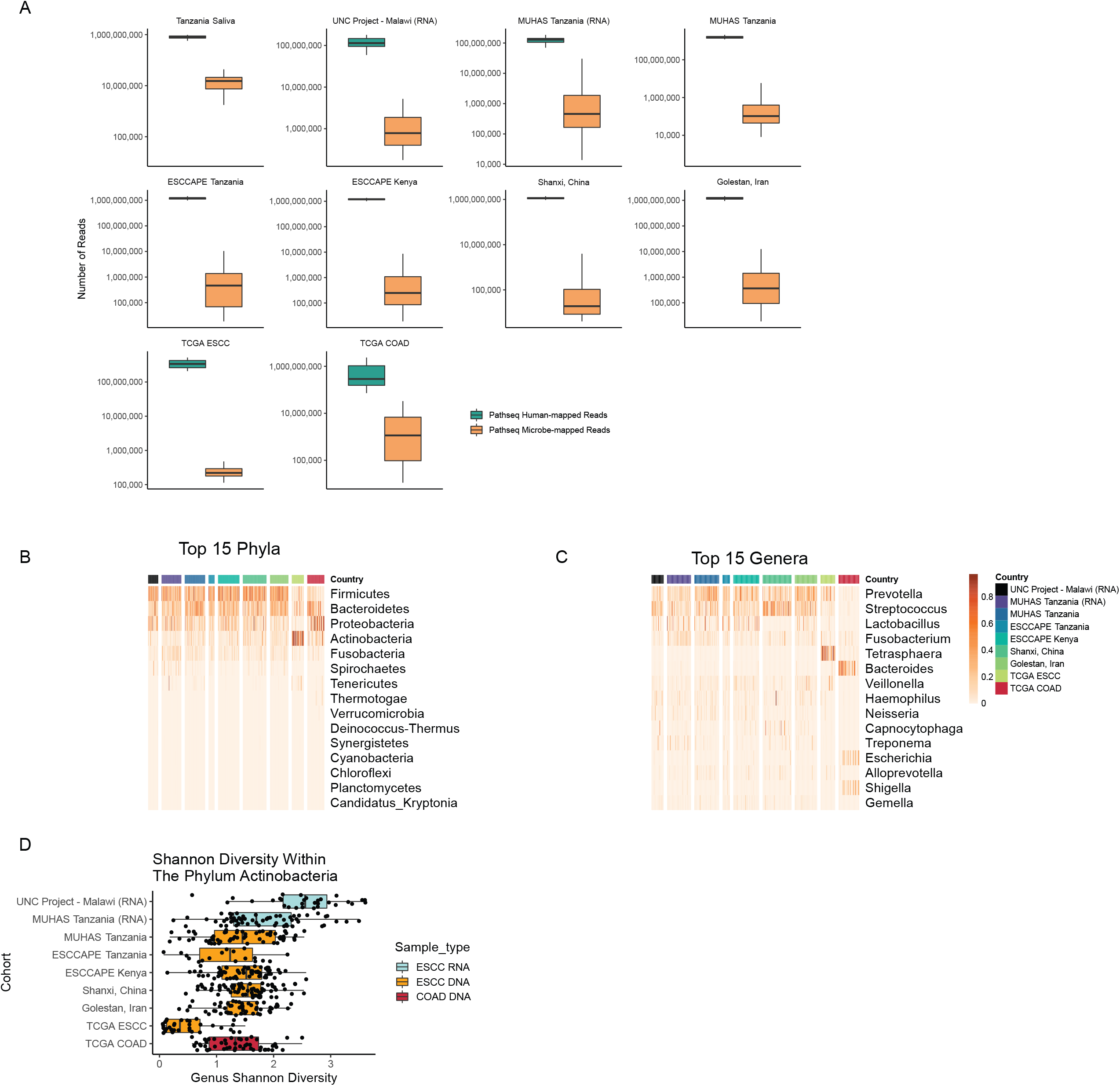
GATK-PathSeq statistics and extended phyla and genera information. A. Boxplots indicating the number of GATK-PathSeq Human-mapped reads and GATK-PathSeq microbe-mapped reads for each patient cohort. Samples from each cohort are WGS unless noted with “(RNA)”, in which case they are RNAseq. B. Heatmap describing the relative abundance of the 15 top phyla sorted by average phylum relative abundance. Each column represents one sample. Rows represent the indicated phyla. Units are relative abundance. Samples from each cohort are WGS unless noted with “(RNA)”, in which case they are RNAseq. C. Heatmap describing the relative abundance of the 15 top genera sorted by average genera relative abundance. Each column represents one sample. Rows represent the indicated genera. Units are relative abundance. Samples from each cohort are WGS unless noted with “(RNA)”, in which case they are RNAseq. D. Boxplot representing the Shannon diversity of genera that fall within the phylum *Actinobacteria* for each patient in each cohort. Samples from each cohort are WGS unless noted with “(RNA)”, in which case they are RNAseq.

**Figure S2.**
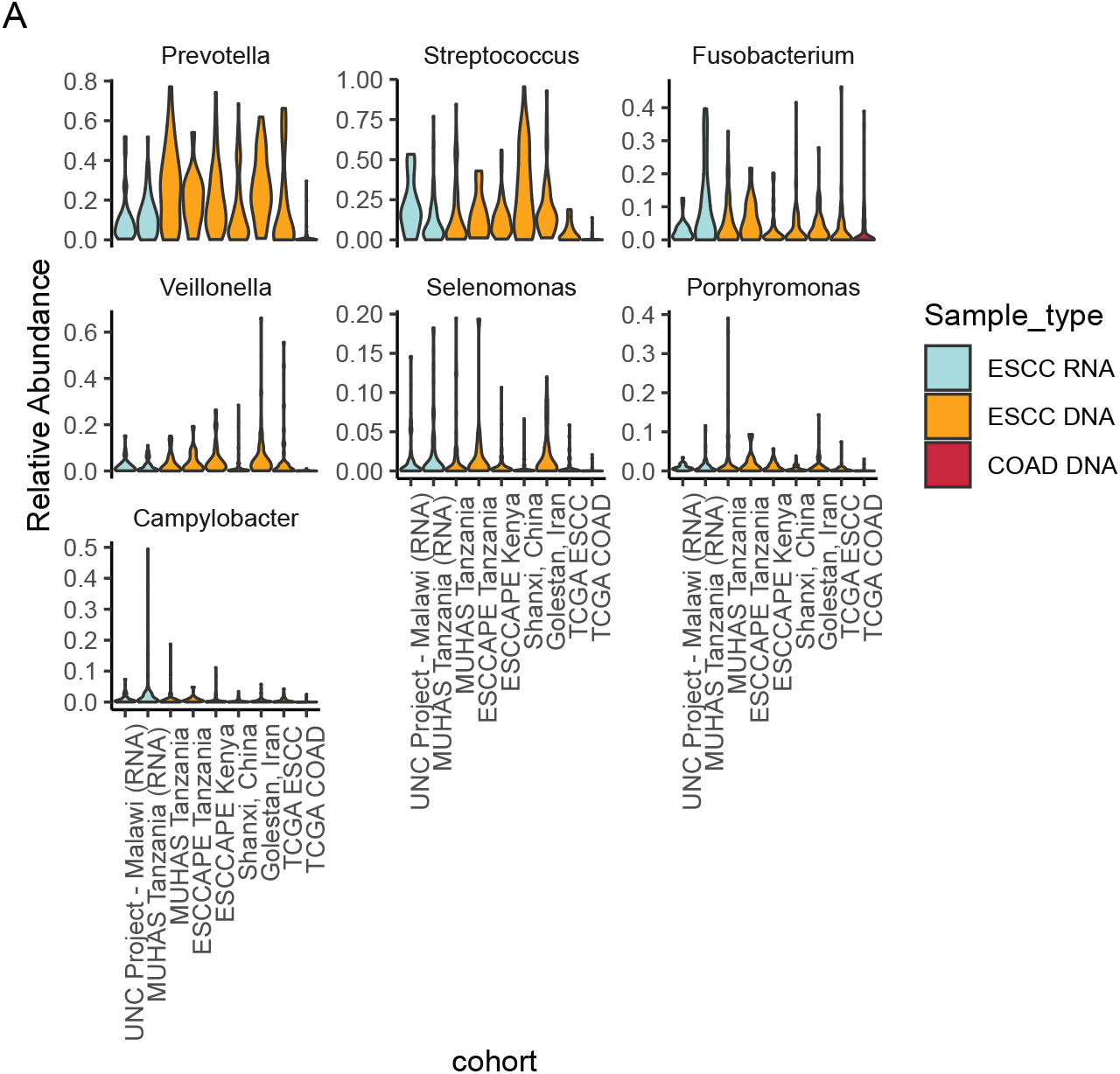
Distribution of *Fusobacterium, Selenomonas, Prevotella, Streptococcus, Porphyromonas, Veillonella*, and *Campylobacter* relative abundance of genus reads for all samples in each study. A. The distribution of the relative abundance of genus-mapping reads for seven selected genera in all studies. The width of each violin represents the proportion of samples which have the indicated relative abundance of each genus. In contrast to **Figure 2D**, which only plots up to 50 samples per study, this plot includes all patients. Samples from each study are WGS unless noted with “(RNA)”, in which case they are RNAseq.

